# Impact of branching on the conformational heterogeneity of the lipopolysaccharide from *Klebsiella pneumoniae*: Implications for vaccine design

**DOI:** 10.1101/500645

**Authors:** Asaminew H. Aytenfisu, Raphael Simon, Alexander D. MacKerell

**Affiliations:** University of Maryland Computer-Aided Drug Design Center, Department of Pharmaceutical Sciences, School of Pharmacy, University of Maryland, Baltimore, Maryland 21201, United States; Center for Vaccine Development, Institute for Global Health, School of Medicine, University of Maryland, Baltimore, Maryland 21201, United States

## Abstract

Resistance of *Klebsiella pneumoniae* (KP) to antibiotics has motivated the development of an efficacious KP human vaccine that would not be subject to antibiotic resistance. *Klebsiella* lipopolysaccharide (LPS) associated O polysaccharide (OPS) types have provoked broad interest as a vaccine antigen as there are only 4 that predominate worldwide (O1, O2a, O3, O5). *Klebsiella* O1 and O2 OPS are polygalactans that share a common D-Gal-I structure, for which a variant D-Gal-III was recently discovered. To understand the potential impact of this variability on antigenicity, a detailed molecular picture of the conformational differences associated with the addition of the D-Gal-III (1→4)Gal branch is presented using enhanced-sampling molecular dynamics simulations. In D-Gal-I two major conformational states are observed while the presence of the 1→4 branch in D-Gal-III resulted in only a single dominant extended state. Stabilization of the more folded states in D-Gal-I is due to a O4-H…O2 hydrogen bond in the linear backbone that cannot occur in D-Gal-III as the O4 is in the Gal*p*(1→4)Gal*p* glycosidic linkage. The impact of branching in D-Gal-III also significantly decreases the accessibility of the monosaccharides in the linear backbone region of D-Gal-I, while the accessibility of the terminal D-Gal-II region of the OPS is not substantially altered. The present results suggest that a vaccine that targets both the D-Gal-I and D-Gal-III LPS can be developed by using D-Gal-III as the antigen combined with cross-reactivity experiments using the Gal-II polysaccharide to assure that this region of the LPS is the primary epitope of the antigen.

**Author Summary:** *Klebsiella pneumoniae* (KP) is a bacterial pathogen commonly associated with hospital acquired infections to antibiotics and is of increasing concern due to the development of resistance to antibiotics including those of last resort. Development of an efficacious KP human vaccine would not be subject to the mechanisms governing antibiotic resistance and, is thus, a public health priority. The present study applies computer simulations to understand the effects of branching on the antigenic D-Gal-I and D-Gal-III O polysaccharide (OPS) species of KP. Results show that branching leads to a single dominate extended conformation, though that conformation occurs in both species. In addition, the terminal D-Gal-II region of both OPS sample similar, exposed conformations while regions of the central repeating units of the OPS are potential hindered from interactions with antibodies in the presence of the branched D-Gal-III OPS. The results suggest a strategy for vaccine development based on D-Gal-III as the antigen combined with cross reactivity experiments to assure that the D-Gal-II region is the primary epitope of the antigen.

## Introduction

*Klebsiella pneumoniae* is a gram negative encapsulated bacterial pathogen, common in the environment, and is a major cause of hospital acquired infections, for which the recent development of widespread antimicrobial resistance among clinical isolates has become an urgent threat.[1] Individuals with impaired host defenses due to chronic illness or immunosenescence are generally at highest risk[2-6] With the limited pipeline of new and novel antibiotics, development of an efficacious *Klebsiella* vaccine is urgently needed. *K. pneumoniae* typically expresses both lipopolysaccharide (LPS), comprised of a conserved core polysaccharide (CP) linked to lipid A and a repeating polymer of O polysaccharide (OPS), and a capsular polysaccharide (CPS, K-antigen), both of which contribute to virulence. While there are greater than 80 different *Klebsiella* capsule serotypes for which no single type predominates, there are only 8 recognized OPS serotypes of which 4 (O1, O2a, O3, O5) account for most human disease globally.[2, 7, 8] Thus, the development of vaccines based on OPS is preferable due to the lower valency requirement to enable broad coverage. However, variability in the composition of OPS subtypes may impact the utility of OPS as a *K. pneumoniae* vaccine antigen.^[2-4, 9-11]^

*Klebsiella* O1, a poly-galactan, is formed by a short stretch of a D-Galactan-I (D-Gal-I) comprised of repeats of ⟶3)-β-Gal*f*-(1⟶3)-α-Gal*p*-(1⟶, that is linked at the non-reducing end to a longer stretch of D-galactan-II repeat units (D-Gal-II) that is generated by ⟶3)-β-Gal*p*-(1⟶3)-α-Gal*p*-(1⟶. The D-Gal-I structure, when uncapped with D-Gal-II, forms the O2a serotype.[4] A recent report documented a variant of D-Gal-I, that contains a (1⟶4)-α-Gal*p* branch, which has been designated as D-Gal-III.[12] In this work, we investigate the conformational properties associated with the addition of this branch to the *K. pneumoniae* O1/O2a OPS (Scheme 1).[4, 12] To investigate the conformational properties we apply computational molecular dynamics (MD) simulations with enhanced sampling via Hamiltonian replica exchange, a technique that has been successfully used to elucidate the conformational properties of other polysaccharides.[3, 4, 13-20] Enhanced sampling was achieved through the use of the Solute Tempering 2 method (HREST) in conjunction with the use of biasing potentials in the context of the correction map (CMAP)[21, 22] approach, termed HREST-bpCMAP.[18, 20, 23-26] This approach is applied in the present study to elucidate the changes in the conformational properties associated with the addition of 1⟶4 branches to the D-Gal-I repeat units (RU) yielding D-Gal-III.[27] The mechanism by which branching on the *K. pneumoniae* serotypes O1 and O2a that may impact antigenicity is explored in terms of both the conformational properties and accessibility of the monosaccharide components of the LPS.

D-Gal-I unit: ⟶3)-β-D-Gal*f*-(1⟶3)-α-D-Gal*p*-(1⟶

D-Gal-III unit: ⟶3)-β-D-Gal*f*-(1⟶3,4)-α-D-Gal*p*-(1⟶[⟶1)-α-D-Gal*p*]

D-Gal-II unit: ⟶3)-β-D-Gal*p*-(1⟶3)-α-D-Gal*p*-(1⟶

Core polysaccharide (CP) -β-GlcNAc(1⟶5)i α -Kdo(2⟶6)-[– α -Hep(1⟶4)-anhMan

D-Gal-I: [⟶3)-β-D-Gal*p*(1⟶3)-α-D-Gal*p*(1⟶3)}_3_{-β-D-Gal/(1⟶3)– α -D-Gal*p*(1⟶3)}_n_-β-GlcNAc(1⟶5)– α -Kdo(2⟶6)-[– α -Hep(1⟶4)]-anhMan

D-Gal-III: ⟶3)-β-D-Gal*p*(1⟶3)– α -D-Gal*p*(l⟶3)}_3_{-β-D-Gaį/(l⟶3)– α -D-Gal*p*(1⟶3)[-α-D– Gal*p*(1⟶4)]}_n_-β-GlcNAc(1⟶5)-α -Kdo(2⟶6)-[– α -Hep(1⟶4)-anhMan

Scheme 1) Sequences of the D-Gal-I, D-Gal-III, D-Gal-II repeating subunits, the core polysaccharide (CP) and of the full D-Gal-I and D-Gal-III polysaccharides considered in the present manuscript where n indicates the number of repeating units (n = 3, 4 or 5).[4, 28]

## Results and Discussion

The goal of the present study was to investigate the impact of the 1⟶4-Gal branch on the conformational properties and accessibilities of the *K. pneumoniae* O1 and O2a OPS using explicit solvent MD simulations (Table 1). As shown in Fig 1 the OPS are comprised of terminal D-Gal-II and outer core polysaccharide (CP) regions linked to D-Gal-I and D-Gal-III RUs for D-Gal-I and D-Gal-III, respectively. The number of RUs was set to 3, 4 or 5 in each OPS structure to determine their potential impact on the overall conformational properties.[27] The change in the number of RUs was found to have minimal impact on the overall conformational properties.

**Table 1).** Systems studied.

| Label |  | Repeating Units (RU) |  |  |  | # of atoms |
| --- | --- | --- | --- | --- | --- | --- |
|  |  | Gal-II | Gal-I | Gal-III | CP |  |
| D-Gal-I | 3RU | 3 | 3 |  | 1 | 30330 |
|  | 4RU | 3 | 4 |  | 1 | 36138 |
|  | 5RU | 3 | 5 |  | 1 | 49797 |
| D-al-III | 3RU | 3 |  | 3 | 1 | 30303 |
|  | 4RU | 3 |  | 4 | 1 | 36168 |
|  | 5RU | 3 |  | 5 | 1 | 49713 |

**Fig 1.**
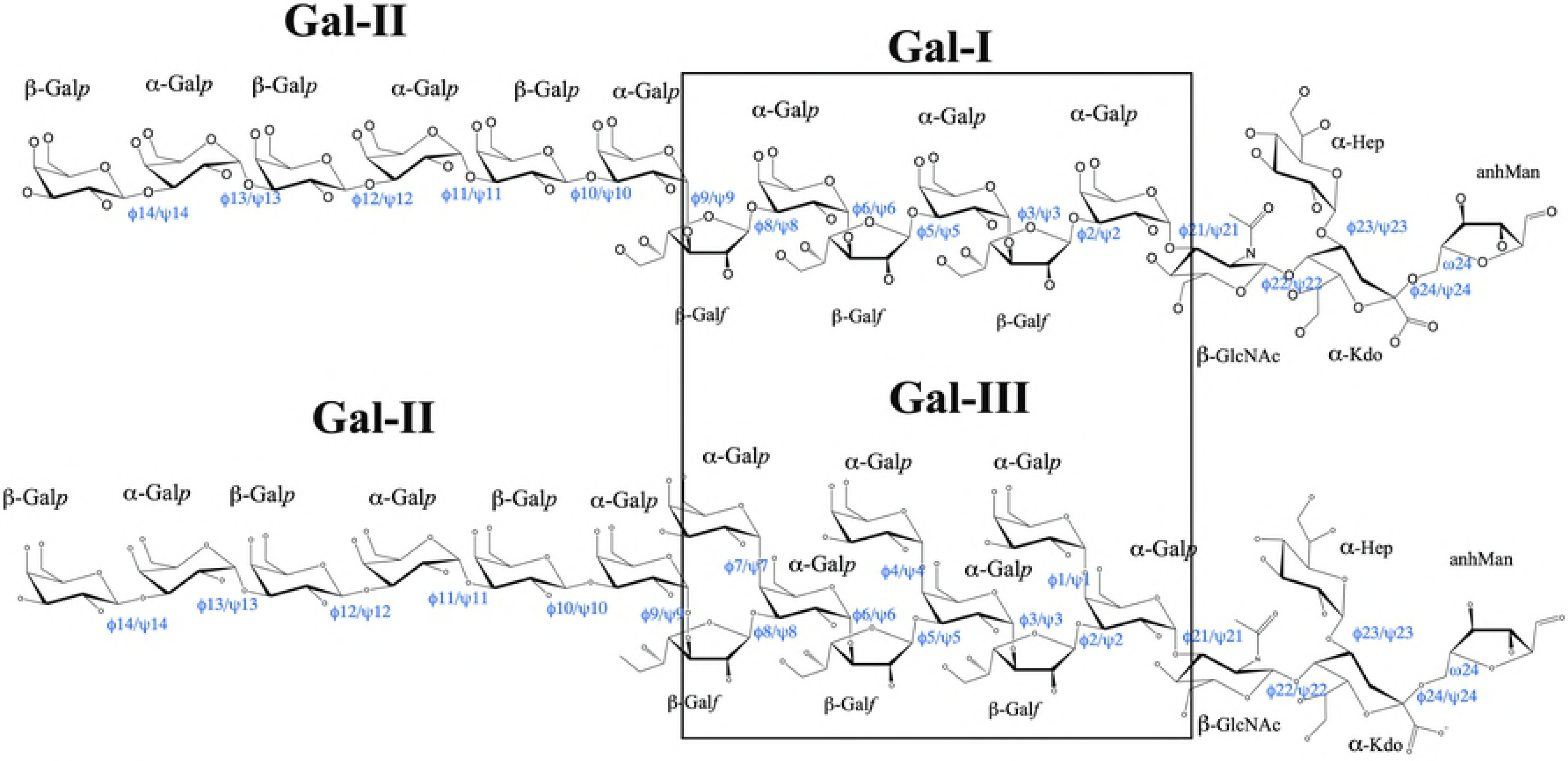
Structures of the **D-Gal-I** (upper) and **D-Gal-III** (lower) polysaccharides. *K. pneumoniae* O1 OPS has an additional D-Gal-II OPS RUs attached to the non-reducing end. The Core Polysaccharide (CP) to the right represents the outer part of the core polysaccharide that would be linked the outer membrane lipid bilayer.

Initial conformational analysis involved the end-to-end distance distributions for the entire OPS and the linear backbone regions of the OPS, with the analysis then progressing to local interactions. As may be seen in Fig 2, there is a systematic increase in the sampling of longer conformations in the D-Gal-III OPS, though both OPS do sample similar fully extended conformations. The trends are consistent across the three numbers of RUs studied. To identify the cause of the conformational difference between D-Gal-I and D-Gal-III, local distance distributions between the RU_n_(C3) to RU_n+1_(C1) atoms in adjacent RUs were determined and presented as probability distributions for each OPS as shown on Fig 3. There are two major populations around 5.2 Å and 6.7 Å for D-Gal-I, while a single population dominates around 6.7 Å for D-Gal-III. This analysis indicates that local conformational differences were occurring between the RUs due to the alteration in branching. The results suggest that there are specific interactions contributing to the different distance distributions as the number of RUs has no significant effect on this local difference between D-Gal-I and D-Gal-III.

**Fig 2.**
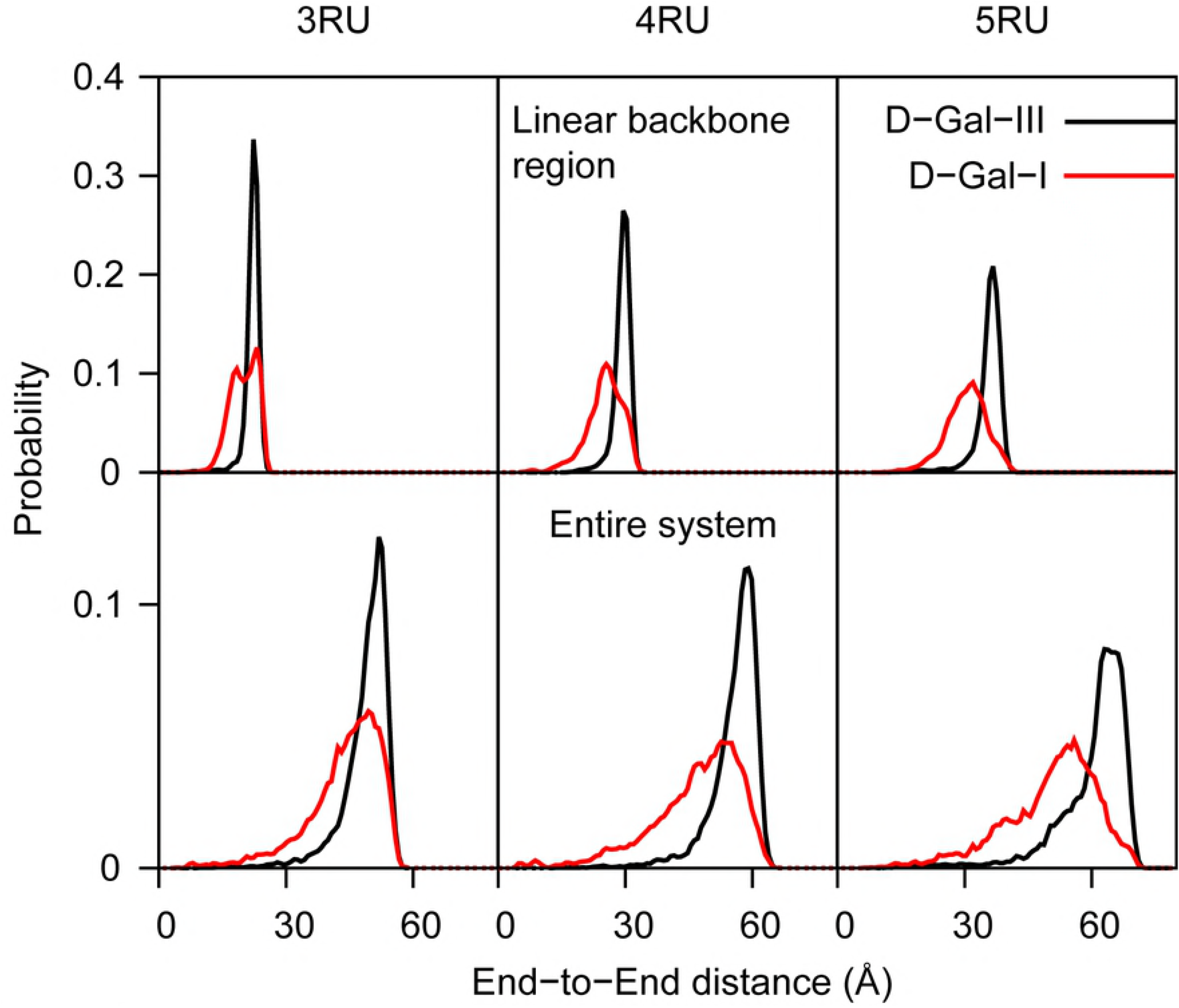
Polysaccharide end-to-end distance distributions based on the CP anhMan C1 atom and the Gal-II terminal β-Gal*p* C1 atom. The distance distributions for the linear backbone region is based on the C1 atoms of the α-Gal*p* of the terminal RU and of the first α-Gal*p* of D-Gal-II. Top and bottom rows correspond to linear backbone region and entire OPS end-to-end distance, respectively, with the black solid curves representing the D-Gal-III and the red dash curves representing the D-Gal-I. Number of Repeating Units 5RU, 4RU and 3RU are plotted along the column from left to right.

**Fig 3.**
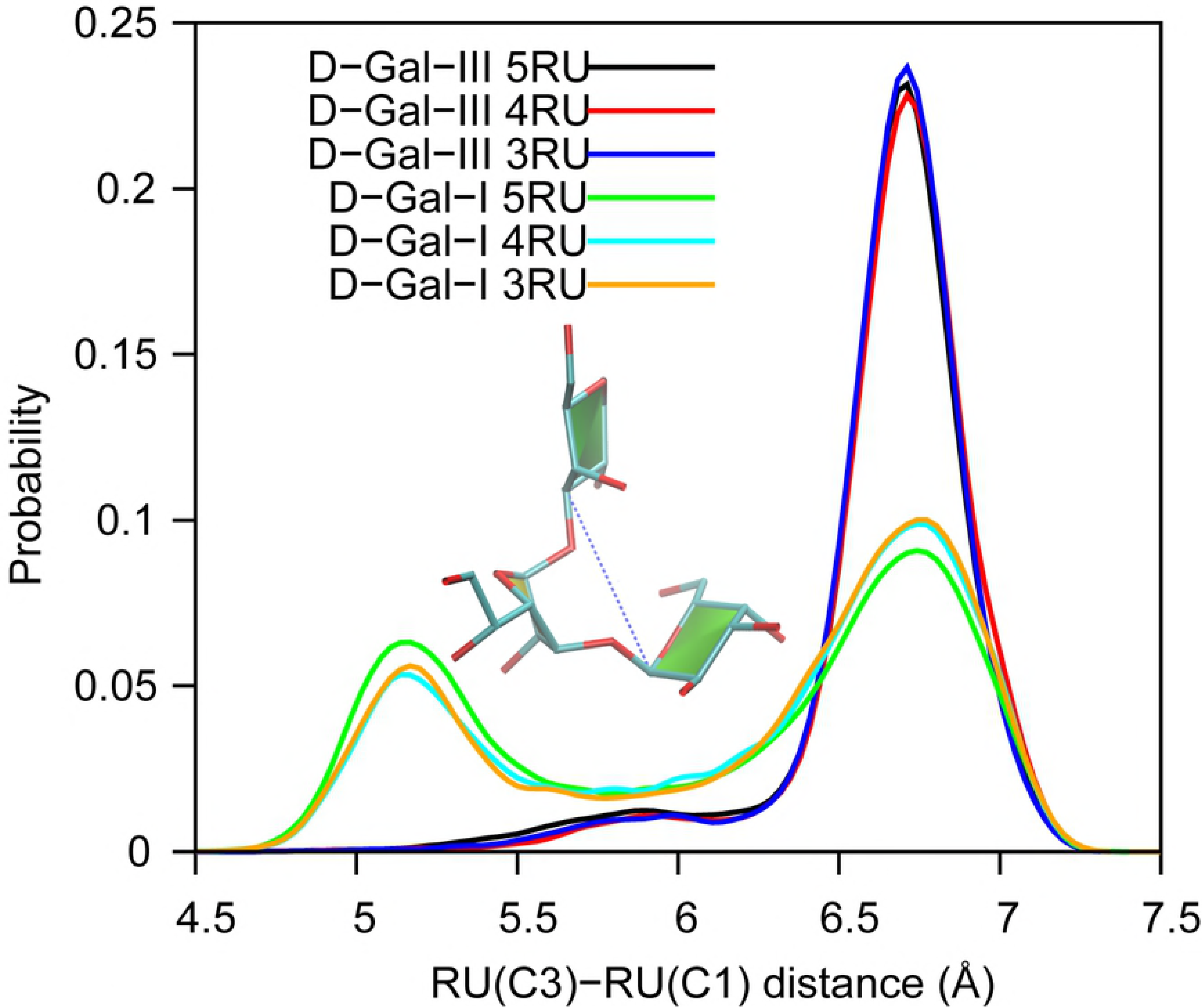
Adjacent RU-RU C1 to C3 distance probability distributions. Distributions include all C1-C3 distances in the linear backbone regions as indicated in the inset structure of a Gal-I RU.

To identify the underlying local interactions that may lead to the conformational difference between D-Gal-I and D-Gal-III, hydrogen bond analysis was performed in the linear backbone region of the OPS. This analysis identified hydrogen bonds involving the O2 and O4 atoms of the α-D-Gal*p* monosaccharides as having significant differences between the two classes of OPS. As shown in Fig 4 the probability distribution of the O4 to O2 distances show distinct differences, with shorter distances sampled in the D-Gal-I OPS. To determine if this interaction was related to the shorter RU-RU distances occurring in D-Gal-I, the correlation of the RU_n_(O4)-RU_n+1_(O2) distance with the RU_n_(C3)-RU_n+1_(C1) distance for the D-Gal-I was analyzed (Fig 5). Analysis of Fig 5 shows the short RU_n_(O4)-RU_n+1_(O2) hydrogen bonds to be highly correlated with the shorter RU_n_(C3)-RU_n+1_(C1) distances, indicating this interaction to lead to the sampling of the shorter RU-RU distance distributions in D-Gal-I. Notably, in D-Gal-III this interaction is not possible due to the O4 atom participating in the glycosidic linkage associated with the additional 1⟶4 branch in D-Gal-III, such the O4 can no longer act as a hydrogen bond donor. An image of the O2-O4 hydrogen bonds occurring in D-Gal-I that leads to the sampling of shorter distances is shown in Fig 5. This behavior is clear when analyzing the specific RU-RU pair distances showing the distinct peaks in Fig 3. However, when analyzing the overall full OPS and linear backbone region end to end distances in Fig 2 distinct peaks are not observed. This is due to the presence of multiple RU in each OPS, yielding different combinations of short and long RU-RU distances for each adjacent RU pair such that broad distributions are observed for the full linear backbone regions. In the case of the 3RU D-Gal-I small peaks are observed for the linear backbone region end to end distance distribution as there are only two RU-RU combinations that can occur.

**Fig 4.**
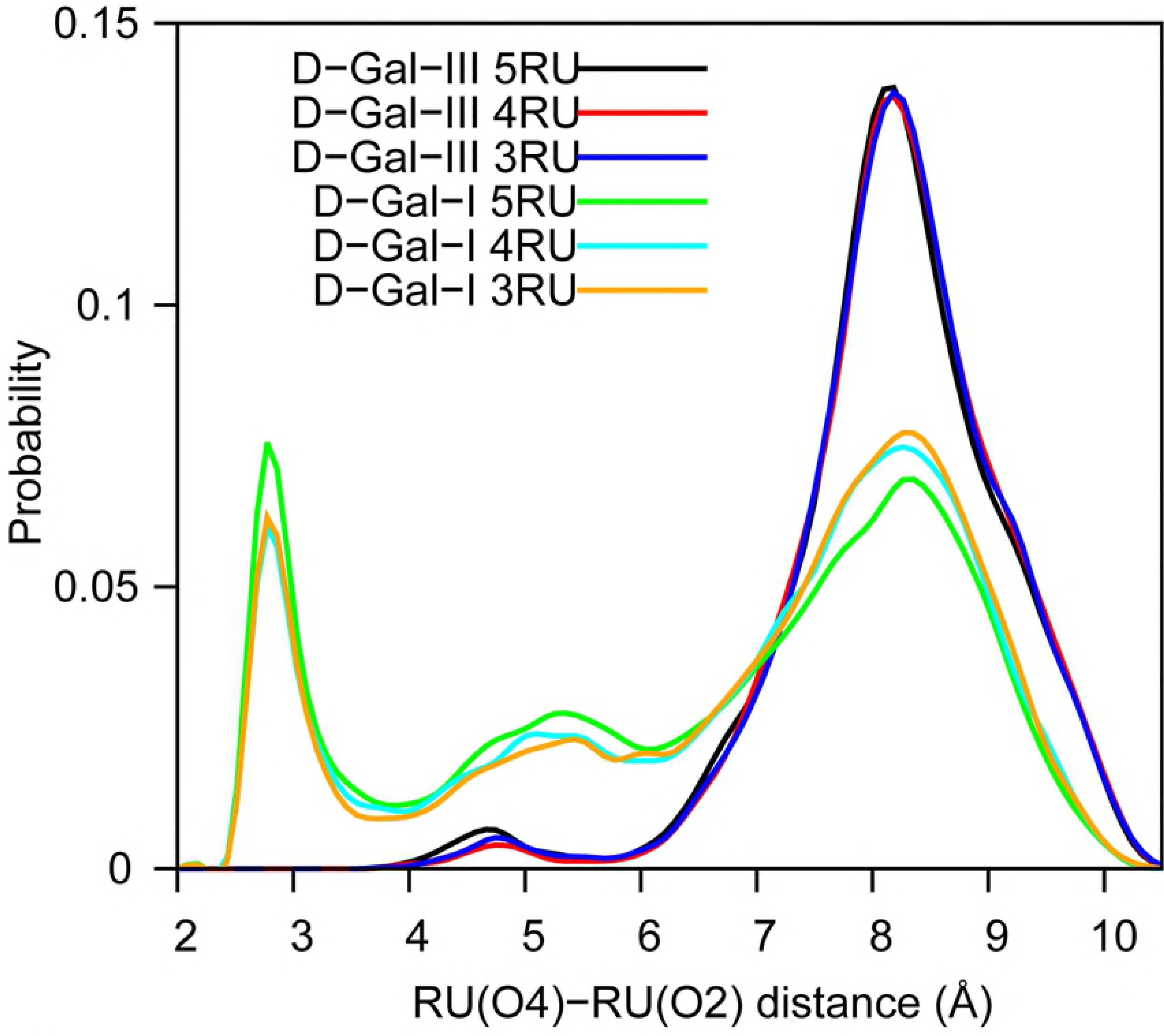
Adjacent RU-RU O2 to O4 distance probability distributions. Distributions include all O2-O4 pairs in the linear backbone region of D-Gal-I and D-Gal-III.

**Fig 5.**
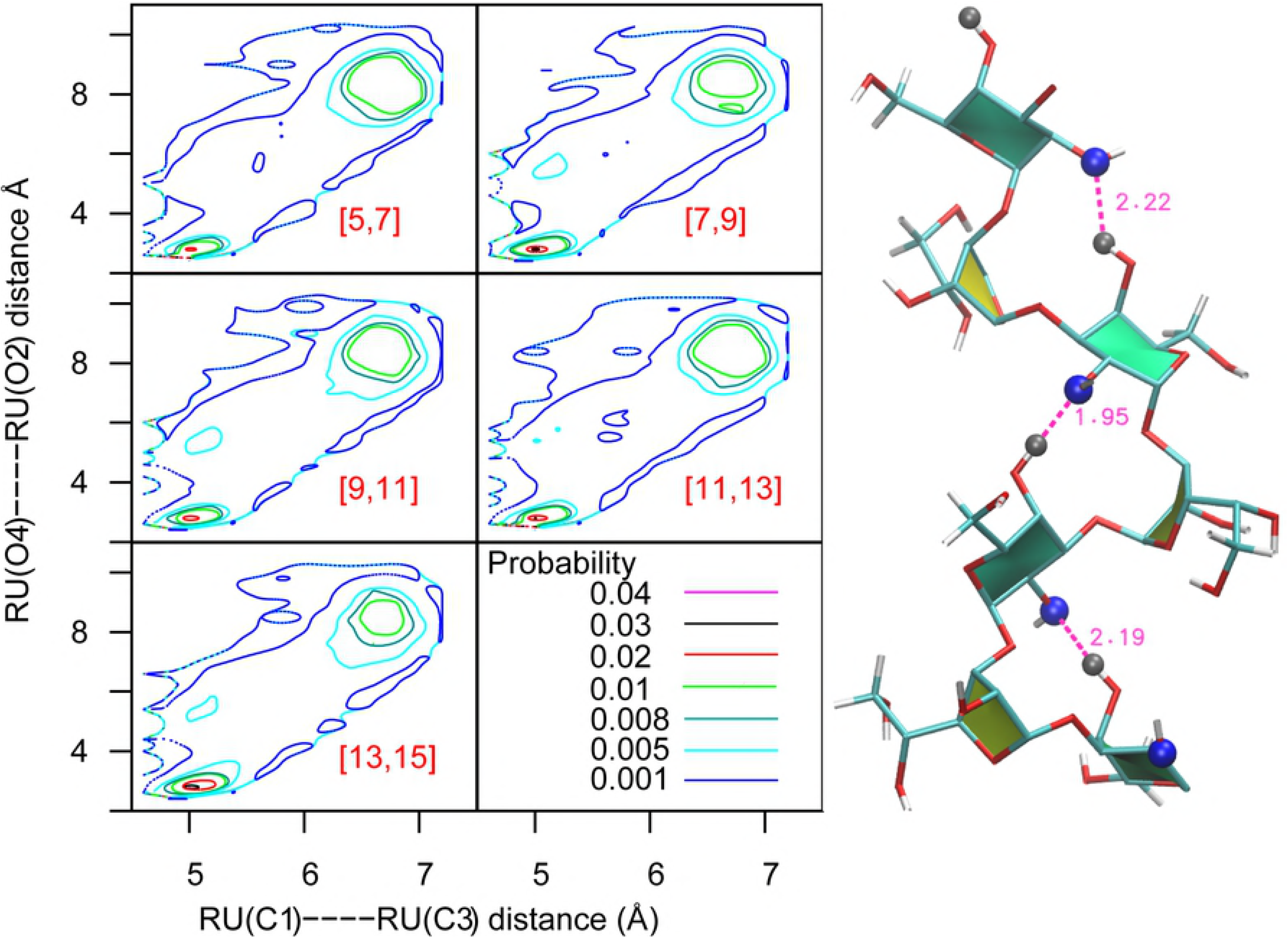
Probability distribution of the linear backbone adjacent RU_n_(O4)-RU_n+1_(O2) versus RU_n_(C1)-RU_n+1_(C3) distances of D-Gal-I. The x-axis corresponds to the linear backbone RU_n_(C1)-RU_n+1_(C3) distances shown on Fig 3 and the y-axis corresponds to the RU_n_(O4)-RU_n+1_(O2) distances shown on Fig 4. The HO4 and O2 atoms form a hydrogen bond is shown in the right panel. The hydrogen bond between HO4 and O2 occurs at shorter RU-to-RU distance, indicating that the missing HO4 from D-Gal-III contributes to folding of the D-Gal-I. For clarity aliphatic hydrogens are omitted and pairs of RUs are labeled in red.

To understand if any additional interactions may be present in D-Gal-III that impact the conformational properties, analysis was performed on interactions involving the D-Gal-III branched α-D-Gal*p* monosaccharides. The prominent hydrogen bond that is present in D-Gal-III is between the O6 atoms of adjacent RUs in the linear backbone region. Plotted in Fig 6 are the probability distributions for the distance between RU_n_(O6(H)) and RU_n+1_(O6(H)). Evident is the presence of high populations of O6-O6 interactions occurring at short distances corresponding to hydrogen bonds, with that high population increased in the 4RU and 5RU species. While this interaction is present to varying degrees in D-Gal-III OPS, as there is only a single distribution of RU(C1)-RU(C3) distances for D-Gal-III, it does not significantly impact the overall conformation of the OPS. However, this interaction may impact the accessibility of the α-D-Gal*p* monosaccharide in the branch, as discussed below.

**Fig 6.**
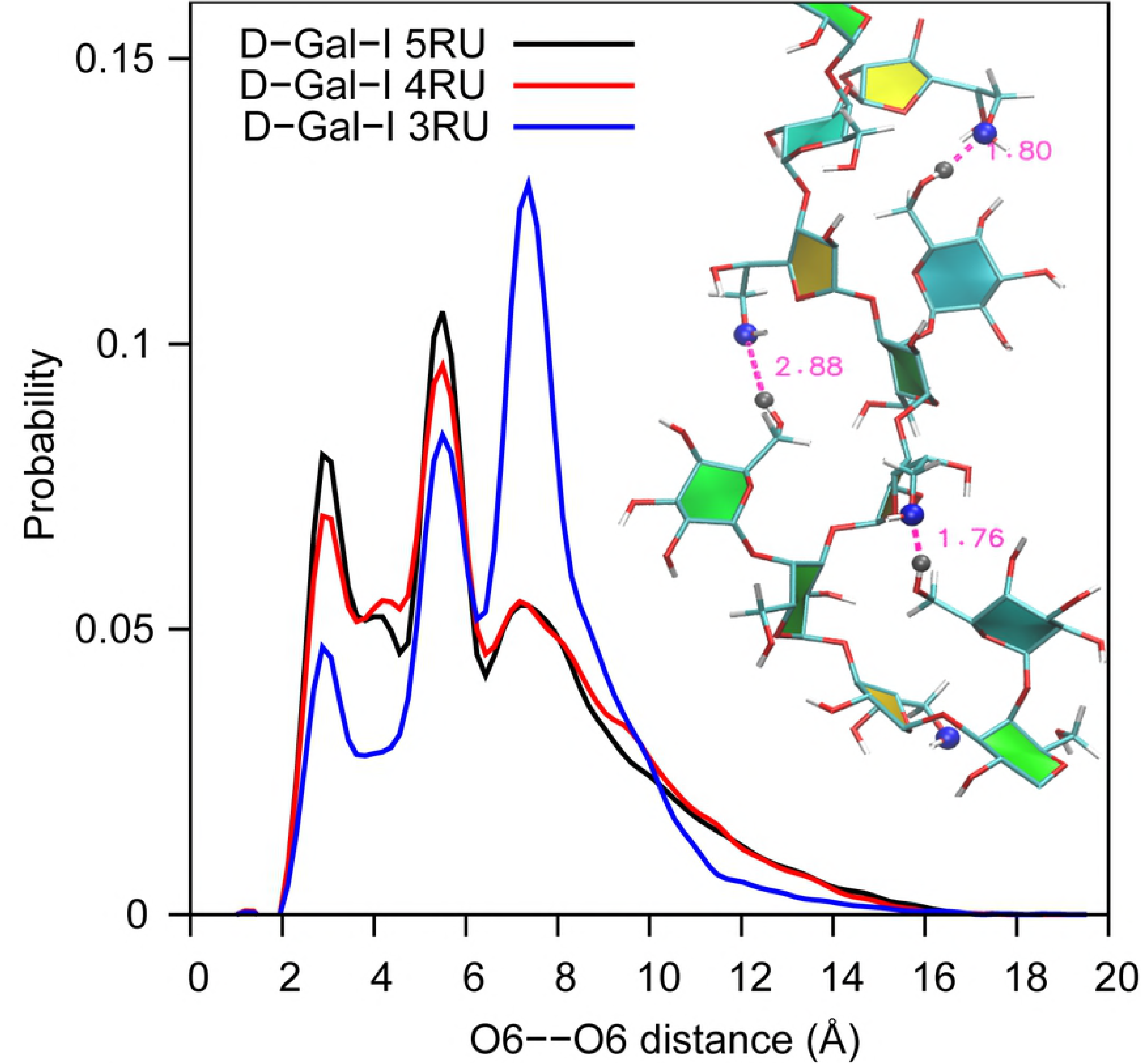
RU to RU O6(H)-O6 distance probability distributions. Distance distributions between O6(H) atoms of the D-Gal-III galactopyranose (Gal*p*) and the galactofuranose (Gal*f*) of the adjacent RU monosaccharides in D-Gal-III. The O6…H-O6 atoms form a hydrogen bond as shown in the inset. There is a preference for hydrogen bonds based on number of RUs. For clarity, aliphatic hydrogens are omitted.

### Antibody Accessible Surface Areas (AASA)

Understanding the regions of the OPS antigens exposed and accessible to the environment indicates those surfaces that can partake in direct interactions with antibodies, thereby contributing to antigenicity. Such information has the potential to further clarify whether the antigenicity of the polysaccharide variants may be similar, thereby facilitating vaccine design. To quantify environmental exposure, we use the AASA, which is analogous to the solvent accessible surface area but calculated with a larger radius for the probe molecule, thereby modeling regions of an antigen that can interact with an antibody. The AASA of the monosaccharides in the D-Gal-I and D-Gal-III OPSs are shown in Fig 7 for the 3, 4 and 5 RU systems. Increasing the number of RUs does not impact the antibody exposure in all of the monosaccharides. However, when comparing the unbranched vs. branched OPS it is evident that the exposure of both the β-D-Gal*f* (Ax in Fig 7) and α-D-Gal*p* (Cx in Fig 7) monosaccharides that are common to both D-Gal-I and D-Gal-III is substantially decreased in the presence of the D-Gal*p* (Dx in Fig 7) branched monosaccharides. This decreased exposure is facilitated by the O6(H) to O6 hydrogen bonds associated with the branch in D-Gal-III discussed above (Fig 6). As expected the D-Gal*p* (Dx in Fig 7) monosaccharides are highly exposed in all cases. Concerning the terminal D-Gal-II and CP regions, the antibody exposure is similar for D-Gal-I and D-Gal-III.

**Fig 7.**
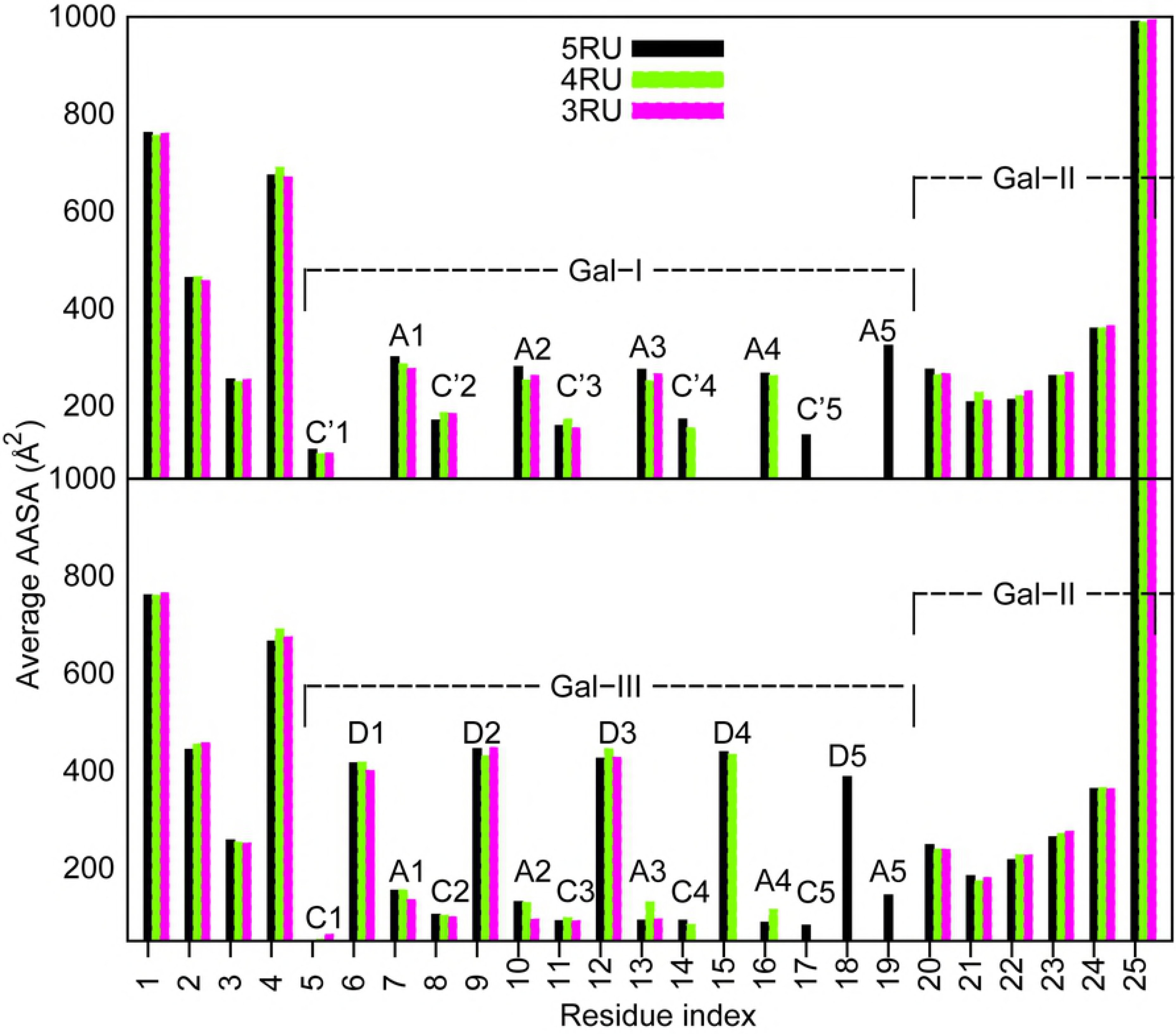
Average Antibody Accessible Surface Area (AASA, Å^2^) for the D-Gal-I (upper panel) and D-Gal-III (lower panel) OPS. Ax indicates β-D-Gal*f* Cx indicates α-D-Gal*p* and Dx indicates D-Gal*p* monosaccharides where x indicates the RU number. Results are averaged over the entire HREST2-bpCMAP trajectories. AASA value is calculated using a probe radius of 10 Å and an accuracy of 0.5.

As both species of OPS can sample the extended states, additional AASA analysis focused on the accessibility of the monosaccharides in only the extended states for both D-Gal-I and D-Gal-III. Direct comparison of the results for the two 5RU species is shown in Fig 8. For the terminal regions the exposure of the monosaccharides is similar; however for the linear backbone region the presence of the branching α-D-Gal*p* moieties (D in Fig 8), which are highly exposed, lead to significant decreases in the exposure of the β-D-Gal*f* and D-Gal*p* monosaccharides (individual AASA for the different number of RUs are included in supporting information Fig S3A-B). These results indicate, assuming the extended conformations participate in interactions with antibodies, that if the interactions are dominated by the terminal D-Gal-II polysaccharide, the presence of the branches in D-Gal-III may not impact antigenicity. However, if the antibody-antigen interactions involve the RUs of the linear backbone regions, those interactions would be significantly altered.

**Fig 8.**
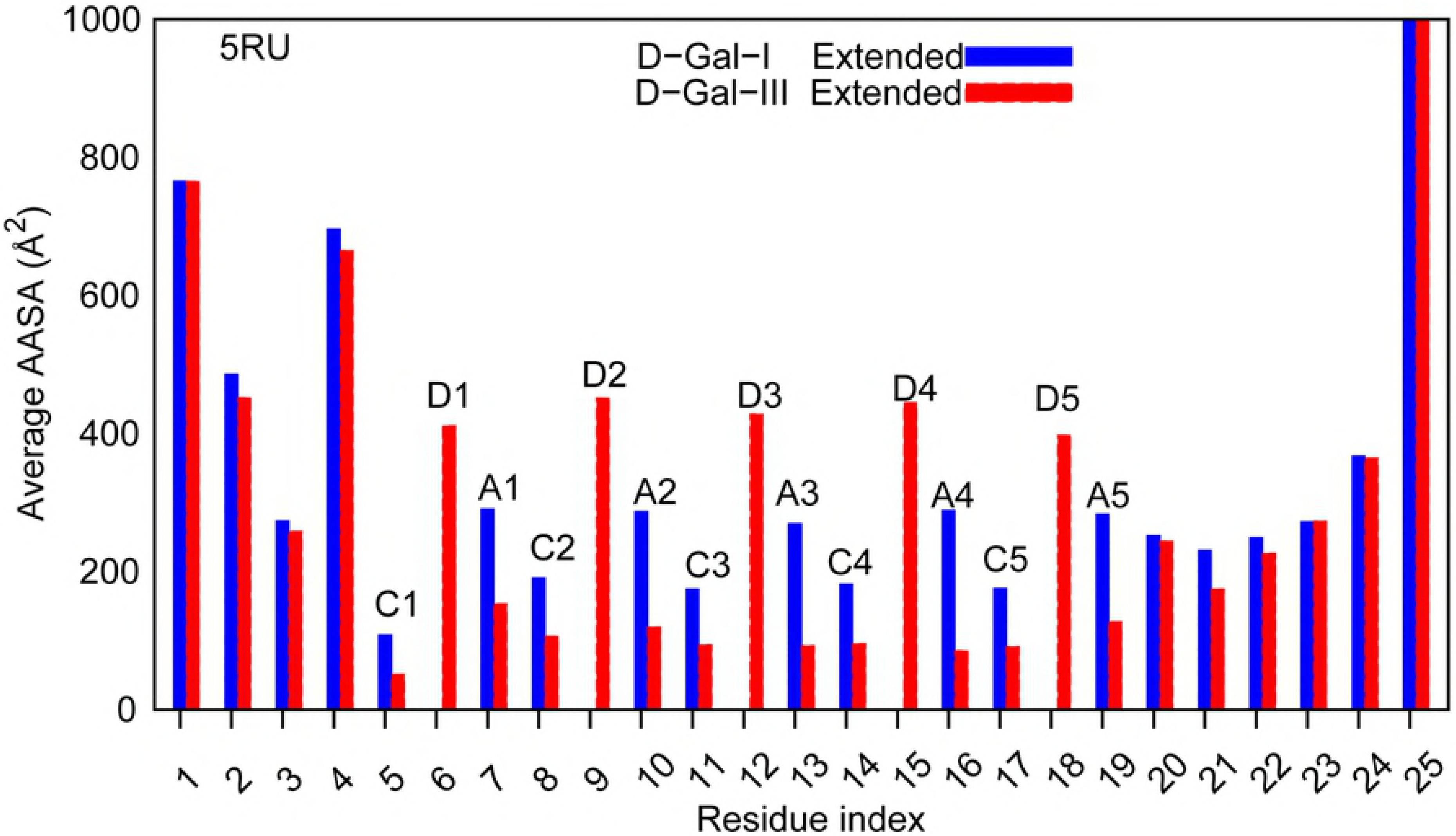
Average Antibody Accessible Surface Area (AASA, Å^2^) for the extended conformations of the monosaccharides in the D-Gal-I (blue) and D-Gal-III (red) OPS for the 5RU species. Extended conformations were based on end to end distances > 60 Å (see Fig 2). Ax indicates β-D-Gal*f* and Dx indicates D-Gal*p* monosaccharides where x indicates the RU number. Results are averaged over the entire HREST2-bpCMAP trajectories.

### Glycosidic linkage-based clustering analysis (GL)

Additional analysis was undertaken to obtain a more quantitative estimate of the conformational states being sampled by the two OPS. Conformations of the saccharides were identified based on clustering of glycosidic linkages (GL) and their sampled populations determined, as previously presented.[18, 23, 24, 46] To perform GL clustering it is necessary to assign the different glycosidic conformations to integers on which the clustering is performed. Shown in Fig S4A-B are 2D φ/ϕ potential of mean force surfaces partitioned into 8 quadrants corresponding to local minima sampled by the Gal*p*(1⟶3)Gal*f* and Gal*f*(1⟶3)Gal*p* glycosidic linkages, respectively. Gal*p*(1⟶3)Gal*f* corresponds to the linkage labeled by φ3/ϕ3, φ6/ϕ6, φ9/ϕ9 and Gal*f*(1⟶3)Gal*p* corresponds to the linkage labeled by φ2/ϕ2, φ5/ϕ5, φ8/ϕ8 in Fig 1. For the Gal*p*(1⟶3)Gal*f* linkage only 0 and 1 quadrants are sampled significantly with a small amount of sampling of quadrant 4 while quadrant 3 dominates with Gal*f*(1⟶3)Gal*p* with some sampling of quadrant 0. All α-D-Gal*p*(1⟶3)β-D-Gal/(1⟶3)Gal*p* linear backbone region linkages are included in the clustering analysis with the 1⟶4 linkages excluded from the clustering as they do not exist in the D-Gal-I. For the 5RU OPS this yields a total of 10 integers defining the OPS conformations, with the top 10 clusters shown in Table 2. With both the unbranched and branched OPS, the 1111133333 cluster dominates, being responsible for 72% of the conformations with D-Gal-III and 30% with D-Gal-I. With D-Gal-I a number of additional conformations are sampled to significant extents, differing from the most sampled conformation due to changes in one of the α-D-Gal*p* (1⟶3) β-D-Gal*f* glycosidic linkages.

Further analysis focused on the extended conformations (Table 3). Based on the end to end distance of the full 5RU OPS, extended conformations of D-Gal-III are sampled 69% versus 19% with D-Gal-I, consistent with results in Fig 2. As expected, the 1111133333 GL cluster dominates the extended conformations, though there is a decrease in the extent of sampling going from 83% with D-Gal-III to 49% with D-Gal-I, indicating a larger contribution of different conformations to the extended states in the unbranched OPS. Thus, in both the unbranched and branched species significant populations of the extended states are sampled, indicating the importance of those conformations being targeted by antibodies, though a wider range of GL conformations are contributing to the extended states in D-Gal-I.

### 3D spatial distribution

To analyze the spatial extent of sampling of the OPS the 3D spatial volume of the sampled conformations as defined by Cartesian coordinates was calculated. Shown in Fig 9 is the extent of volume sampled at the different isovalues for the D-Gal-I and D-Gal-III 5RU species. The extent of sampling by both types of OPS is large, with the D-Gal-III OPS sampling longer distances, consistent with the analyses above. It is noted that while only a relatively small number of GL clusters are sampled in the simulations (Table 2), the OPS can still sample a wide range of 3D conformational space. However, on the crowded surface of the bacterium the range of sampling may be significantly altered due to interactions with the surrounding environment including adjacent LPS molecules and membrane proteins.

**Fig 9.**
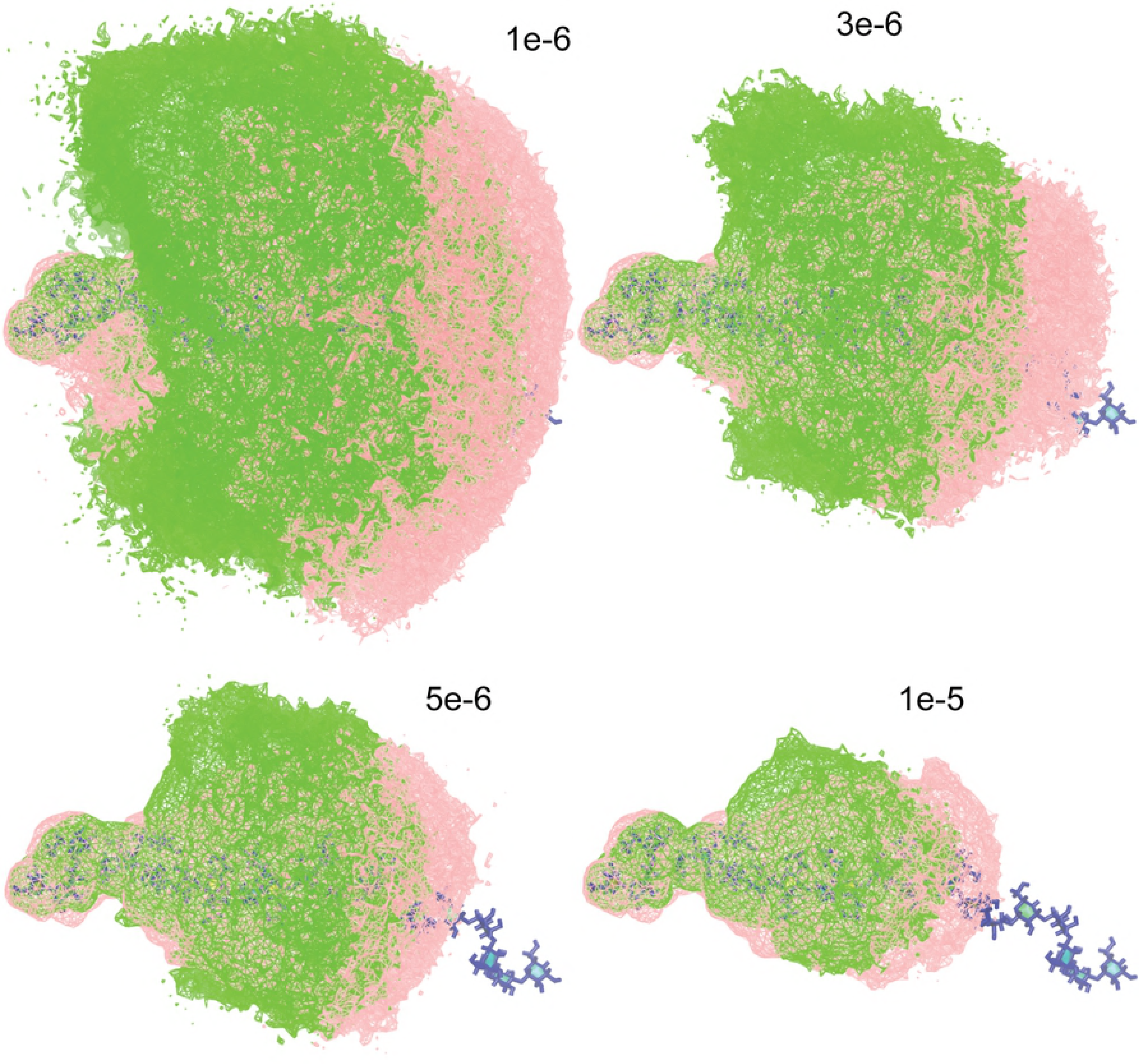
Spatial distribution of 5RU D-Gal-I (green) and D-Gal-III (pink). To compare the extent of conformational sampling between D-Gal-I and D-Gal-III polysaccharides, the 3D spatial distribution (wire frame) are plotted excluding the 1⟶4 branch from the D-Gal-III. The trajectory was aligned with respect to the common part (CP) of the OPS (see Fig 1). The different labels indicate the contour level used to calculate the isosurface in the VMD analysis tool[45], where the voxels were normalized with respect to the total number of voxels sampled such that their sum equals 1.

## Conclusion

The presented analysis yields a detailed molecular picture into the conformational differences between D-Gal-I and D-Gal-III Klebsiella O polysaccharides. To understand the effects of branching, MD simulations were undertaken on model D-Gal-I and D-Gal-III species for three different systems with 3, 4 or 5 RUs. The results indicated that the absence of the 1⟶4 branch in D-Gal-III resulted in two dominant conformational states with one compact and a second more extended conformation in the linear backbone region while the presence of this 1⟶4 α-D-Gal*p* branch in D-Gal-III resulted in only a single dominant extended state in the linear backbone region. Stabilization of the compact states in D-Gal-I is due to the O4 – O2 hydrogen bond between adjacent RUs where O4 is a hydrogen bond donor. This interaction cannot occur in the D-Gal-III as the α-D-Gal*p* O4 participates in the (1⟶4) glycosidic linkage to the branched α-D-Gal*p* and can no longer act as a hydrogen bond donor. An additional hydrogen bond, namely between O6 of two consecutive RUs occurs in the linear backbone region of D-Gal-III in the extended state. However, this doesn’t significantly impact the overall conformation of the D-Gal-III as the longer conformation dominates, though it does facilitate shielding of the β-D-Gal*f* and α-D-Gal*p* linear backbone monosaccharides from putative interactions with antibodies.

The present study provides several important insights regarding the use of KP OPS as vaccine immunogens. Monospecific antibodies recognizing D-Gal-III have previously been reported.[3] However, it is unclear whether these monospecific antibodies represent a significant fraction of the complete polyclonal response to the OPS molecule. It is conceivable that cross-reactive antibodies to shared epitopes may be induced following immunization with D-Gal-I/D-Gal-III based vaccines. For a vaccine to successfully target both D-Gal-I and D-Gal-III, our results suggest that it would be advantageous to induce immune responses targeting the extended conformation of the OPS, given the dominance of this latter conformation in D-Gal-III. While a lower population of the extended state is sampled in the D-Gal-I OPS, a significant population of those states (19%) is nevertheless sampled, that should theoretically allow for induction of antibodies targeting this conformation. In addition, it may be anticipated that the process of conformational selection further facilitates the sampling of the appropriate conformation when interacting with such an antibody, analogous to those previously elucidated for HIV envelope glycans.[23] In addition, it has been shown that antibody binding can alter the conformations sampled by an oligosaccharide, which may further facilitate binding.[24, 46]

For an O1 OPS vaccine, despite the reduced accessibility of the β-D-Gal/ and α-D-Gal*p* linear backbone monosaccharides in the RUs of D-Gal-III OPS, antibodies induced against the terminal Gal-II region should be similarly efficacious in the context of D-Dal-I or D-Gal-III given the relative insensitivity of D-Gal-II to the presence of the branches. In addition, as the Gal-II region is membrane distal, it is expected that it would be more exposed compared to the membrane proximal D-Gal-I/D-Gal-III in O1. Indeed, D-Gal-II has been found to represent the primary protective OPS epitope in O1 OPS expressing *Klebsiella*.[3] Furthermore, this epitope has been found to be immunodominant after immunization. Future studies will be needed to characterize the cross-reactive and antigen specific immune responses to O1 and O2 OPS molecules containing D-Gal-I or D-Gal-III.

## Methods

Modeling and simulations were performed with the program CHARMM using the CHARMM36 additive force field for carbohydrates[29, 30] and the CHARMM TIP3P[31] water model. Initial coordinates of the OPS were generated from the topology information present in the force field followed by minimization using the Steepest Descent (SD) and Adopted-Basis Newton-Raphson (ABNR) minimizers for 5000 steps each with end to end distance restraints on the OPS in order to maintain extended conformations. The end to end distance restraint was placed on the anhMan C1 and the terminal β-Gal*p* C1 atoms with a force constant of 15 kcal/mol/Å and with equilibrium distances of 50, 60 and 70 Å for the 3, 4 and 5 RU systems, respectively. The resulting geometries of the OPS were then immersed in a pre-equilibrated cubic water box. The size of the water box was selected based on the condition that it extend at least 10 Å beyond the non-hydrogen atoms of the fully-extended OPS. Water molecules with the oxygen within a distance of 2.8 Å of the non-hydrogen solute atoms were deleted. For all of the subsequent minimizations and MD simulations, periodic boundary conditions were employed using the CRYSTAL module implemented in the CHARMM program.[32-34] A list of the systems studied is shown in Table 1.

Equilibration of the solvated systems was initiated with a 500 step SD minimization followed by a 500 step ABNR minimization in which mass-weighted harmonic restraints of 1.0 kcal/mol/Å were applied on the nonhydrogen atoms of the OPS. Following minimization, each solvated system was initially heated from 100 to 298 K under constant volume and temperature (NVT) followed by 100 ps constant pressure and temperature (NPT) MD simulations at 298 K and 1 atm. In all simulations under the NVT or NPT ensembles, including the subsequent HREST-bpCMAP simulations, the temperature was maintained at 298 K using the Hoover algorithm with a thermal piston mass of 1000 kcal/molps^2^.[35] A constant pressure of 1 atm was maintained using the Langevin piston algorithm with a collision frequency of 20 ps^-1^ and mass of 1630 amu.[36] The covalent bonds involving hydrogen atoms were constrained with the SHAKE algorithm, and a time step of 2 fs was used.[37] In the energy and force evaluations, the nonbonded Lennard-Jones interactions were computed with a cutoff of 12 Å with a switching function applied over the range from 10 to 12 Å. The electrostatic interactions were treated by the particle mesh Ewald method with a real space cutoff of 14 Å, a charge grid of 1 Å, a kappa of 0.34, and the 6-th order spline function for mesh interpolation.[38]

Conformational sampling was enhanced by applying the HREST-bpCMAP method that involves concurrent solute scaling and biasing potentials.[20] All the production HREST-bpCMAP simulations were carried out in CHARMM using the replica exchange module REPDST, with BLOCK to scale the solute-solute and solute-solvent interactions[31, 39, 40] and with specific bpCMAPs applied as the 2D biasing potentials along selected glycosidic linkages.[22] 5RU, 4RU and 3RU (RU=Repeating Unit) OPS were simulated for both D-Gal-I and D-Gal-III, yielding a total of 6 molecules. The RU that corresponds the linear backbone region contains two monosaccharides per RU for the D-Gal-I and three monosaccharide per RU for the D-Gal-III (Scheme 1), while D-Gal-II contains three RU with two monosaccharide per each RU for all 6 systems. The bpCMAP[20] biasing potential is applied to the glycosidic linkages across monosaccharides 5 to 25 for 5RU yielding 20 bpCMAPs for D-Gal-III and 16 bpCMAPs for the D-Gal-I. The biasing potentials are applied for 4RU and 3RU in a similar fashion. The 2-dimensional grid-based bpCMAP were constructed using the corresponding disaccharide model in the gas phase as described previously[18, 20, 41] and was applied along the φ(O_5_-C_1_-O_n_ C_n_)/ϕ(C_1_-O_n_-C_n_-C_n-1_) dihedrals for each glycosidic linkage in the OPS. A total of 8 replicas were carried out for each system and exchanges attempted every 1000 MD steps according to the Metropolis criterion. In HREST-bpCMAP simulations, the solute scaling temperatures were assigned to 298 K, 308 K, 322 K, 336 K, 352 K, 370 K, 386 K and 405 K, with the ground-state replica temperature of 298 K selected to correspond to the experimental studies. The HREST-bpCMAP simulations were run for 150 ns on each OPS/RU combination.

The distribution of scaling factors for the bpCMAPs across the 8 perturbed replicas was determined as previously described and the acceptance ratio between different neighboring replicas was examined to guarantee that sufficient exchanges were being obtained.[18, 20, 23, 24] Analysis of the replica walk for the D-Gal-III ground state replica (Fig S1) showed that multiple transitions from the ground state to the highest replica occurred indicating adequate acceptance rates between the replicas and that the full replica walks from the ground state to highest replica were occurring. Analysis of convergence involved examination of the global and linear backbone end-to-end distance probability distributions from the 0-50, 50-100 and 100-150 portions of the simulations (Supplement information: Fig S2). Final results are presented based on the full 150 ns of sampling obtained in the ground state, 298 K replica.

Antibody accessible surface area (AASA) was calculated using the surf tools in CHARMM by probing with a sphere radius of 10 Å,[33, 42-44] yielding the average AASA of the for the entire OPS as well as for the individual monosaccharides over the entire simulations. Use of AASA allows for prediction of regions of the OPS accessible to direct interactions with antibodies. The AASA is determined in a manner analogous to the solvent accessible surface area calculations and uses a probe radius of 10 Å to approximate an antibody combining region interacting with the antigen.[43, 44]

The OPS conformational sampling in Cartesian space was computed based on the sampled spatial volumes. To compute the sampled spatial volumes (Cartesian space), a 3D grid with a voxel size of 1Å×1Å×1Å was constructed around each saccharide. Then, for a given snapshot from the individual simulations, each voxel was assigned a value of 1 if an atom occupied it with the summation performed over all the snapshots of the trajectories. Final results are normalized for the number of snapshots and the total number of voxels sampled such that the sum of the final voxel occupancies equals 1. Trajectories were aligned to the common terminal monosaccharide. 3D spatial distribution representations were generated using the VMD[45] isosurface tool at isovalues of 1e-6, 3e-6, 5e-6, and 1e-5.

## Acknowledgements

Computational support from the University of Maryland Computer-Aided Drug Design Center is acknowledged.

## Author Contributions

ADM and RS conceived the work. AA performed the calculations. AA and ADM performed the analysis. AA, RS, and ADM wrote and revised the manuscript.

## Supporting Information Captions

Fig S1. Random walk of replica Ground (0) in the space of effective temperature. In this replica walk, D-Gal-I with 4RU is plotted as an example. The walk of the ground state from bottom unperturbed to the highest energy state and down to the ground state indicates efficient conformational sampling.(TIFF)

Fig S2 Convergence of the end-to-end distance based on the 0-50ns, 50-100ns and 100-150ns portions of the enhanced sampling simulations. Top and bottom rows correspond to the entire system and the linear backbone end-to-end distances, respectively. Dash lines correspond to the unbranched and solid lines to the branched OPS. (TIFF)

Fig S3A. Average Antibody Accessible Surface Area (AASA, Å^2^) in compact and extended RU conformations and over the entire D-GAL-I OPS. Results are averaged over the full HREST2-bpCMAP trajectories. AASA value is calculated using a probe radius of 10.0 Å and an accuracy of 0.5. (TIFF)

Fig S3B. Average Antibody Accessible Surface Area (AASA, Å^2^) in compact and extended RU conformations and over the entire D-Gal-III OPS. Results are averaged over the entire HREST2-bpCMAP trajectories. AASA value is calculated using a probe radius of 10.0 Å and an accuracy of 0.5. (TIFF)

Fig S4A. Index of free energy minima of the Gal*p*(1⟶3)Gal*f* glycosidic linkages used in the GL clustering. Torsion angles are given in degrees. In the index table this corresponds to Gal*p*(1⟶3)Gal*f* (TIFF)

Fig S4B. Index of free energy minima of the Gal*f*(1⟶3)Gal*p* glycosidic linkages used in the GL clustering. Torsion angles are given in degrees. In the index table this corresponds to Gal*f*(1⟶3)Gal*p*. (TIFF)

## References

1. Prestinaci F, Pezzotti P, Pantosti A. Antimicrobial resistance: a global multifaceted phenomenon. Pathog Glob Health. 2015;109(7):309–18. https://doi.org/10.1179/2047773215Y.0000000030 PMID:26343252

2. Follador R, Heinz E, Wyres KL, Ellington MJ, Kowarik M, Holt KE, et al. The diversity of Klebsiella pneumoniae surface polysaccharides. Microb Genom. 2016;2(8):e000073. https://doi.org/10.1099/mgen.0O00073 PMID:28348868

3. Stojkovic K, Szijártó V, Kaszowska M, Niedziela T, Hartl K, Nagy G, et al. Identification of D-galactan-III as part of the lipopolysaccharide of Klebsiella pneumoniae serotype o1. Front Microbiol. 2017. https://doi.org/10.3389/fmicb.2017.00684

4. Vinogradov E, Frirdich E, MacLean LL, Perry MB, Petersen BO, Duus J, et al. Structures of lipopolysaccharides from Klebsiella pneumoniae: Elucidation of the structure of the linkage region between core and polysaccharide O chain and identification of the residues at the non-reducing termini of the O chains. J Biol Chem. 2002. https://doi.org/10.1074/jbc.M202683200

5. Broberg CA, Palacios M, Miller VL. Klebsiella: a long way to go towards understanding this enigmatic jet-setter. F1000Prime Rep. 2014;6:64. https://doi.org/10.12703/P6-64 PMID:25165563

6. Podschun R, Ullmann U. Klebsiella spp. as nosocomial pathogens: epidemiology, taxonomy, typing methods, and pathogenicity factors. Clin Microbiol Rev. 1998;11(4):589–603. PMID:9767057

7. Trautmann M, Ruhnke M, Rukavina T, Held TK, Cross AS, Marre R, et al. O-antigen seroepidemiology of Klebsiella clinical isolates and implications for immunoprophylaxis of Klebsiella infections. Clin Diagn Lab Immunol. 1997;4(5):550–5. PMID:9302204

8. Trautmann M, Held TK, Cross AS. O antigen seroepidemiology of Klebsiella clinical isolates and implications for immunoprophylaxis of Klebsiella infections. Vaccine. 2004;22(7):818–21. https://doi.org/10.1016/j.vaccine.2003.11.026 PMID:15040933

9. Ahmad TA, Haroun M, Hussein AA, El Ashry el SH, El-Sayed LH. Development of a new trend conjugate vaccine for the prevention of Klebsiella pneumoniae. Infect Dis Rep. 2012;4(2):e33. https://doi.org/10.4081/idr.2012.e33 PMID:24470947

10. Szijarto V, Guachalla LM, Hartl K, Varga C, Banerjee P, Stojkovic K, et al. Both clades of the epidemic KPC-producing Klebsiella pneumoniae clone ST258 share a modified galactan O-antigen type. Int J Med Microbiol. 2016;306(2):89–98. https://doi.org/10.1016/j.ijmm.2015.12.002 PMID:26723873

11. Whitfield C, Perry MB, MacLean LL, Yu SH. Structural analysis of the O-antigen side chain polysaccharides in the lipopolysaccharides of Klebsiella serotypes O2(2a), O2(2a,2b), and O2(2a,2c). J Bacteriol. 1992;174(15):4913–9. PMID:1378428

12. Stojkovic K, Szijarto V, Kaszowska M, Niedziela T, Hartl K, Nagy G, et al. Identification of d-Galactan-III As Part of the Lipopolysaccharide of Klebsiella pneumoniae Serotype O1. Front Microbiol. 2017;8:684. https://doi.org/10.3389/fmicb.2017.00684 PMID:28487676

13. Kadirvelraj R, Gonzalez-Outeiriño J, Foley BL, Beckham ML, Jennings HJ, Foote S, et al. Understanding the bacterial polysaccharide antigenicity of Streptococcus agalactiae versus Streptococcus pneumoniae. Proc Natl Acad Sci U S A. 2006;103(21):8149.

14. Germer A, Peter MG, Kleinpeter E. Solution-State Conformational Study of the Hevamine Inhibitor Allosamidin and Six Potential Inhibitor Analogues by NMR Spectroscopy and Molecular Modeling. J Org Chem. 2002;67(18):6328–38. https://doi.org/10.1021/jo0163703

15. Fadda E, Woods RJ. Molecular simulations of carbohydrates and protein-carbohydrate interactions: motivation, issues and prospects. Drug Discovery Today. 2010;15(15):596–609. https://doi.org/https://doi.org/10.1016/j.drudis.2010.06.001

16. Wood NT, Fadda E, Davis R, Grant OC, Martin JC, Woods RJ, et al. The influence of N-linked glycans on the molecular dynamics of the HIV-1 gp120 V3 loop. PLoS One. 2013;8(11):e80301. https://doi.org/10.1371/journal.pone.0080301 PMID:24303005

17. Kuttel MM, Jackson GE, Mafata M, Ravenscroft N. Capsular polysaccharide conformations in pneumococcal serotypes 19F and 19A. Carbohydr Res. 2015;406:27–33. https://doi.org/10.1016/j.carres.2014.12.013 PMID:25658063

18. Yang M, Angles D’Ortoli T, Säwén E, Jana M, Widmalm G, Mackerell AD. Delineating the conformational flexibility of trisaccharides from NMR spectroscopy experiments and computer simulations. Phys Chem Chem Phys. 2016. https://doi.org/10.1039/c6cp02970a

19. Kuttel MM, Timol Z, Ravenscroft N. Cross-protection in Neisseria meningitidis serogroups Y and W polysaccharides: A comparative conformational analysis. Carbohydr Res. 2017;446-447:40–7. https://doi.org/10.1016/j.carres.2017.05.004

20. Yang M, Huang J, MacKerell AD, Jr. Enhanced conformational sampling using replica exchange with concurrent solute scaling and hamiltonian biasing realized in one dimension. J Chem Theory Comput. 2015;11(6):2855–67. https://doi.org/10.1021/acs.jctc.5b00243 PMID:26082676

21. MacKerell AD, Feig M, Brooks CL. Improved Treatment of the Protein Backbone in Empirical Force Fields. J Am Chem Soc. 2004. https://doi.org/10.1021/ja036959e

22. MacKerell AD, Feig M, Brooks CL. Extending the treatment of backbone energetics in protein force fields: Limitations of gas-phase quantum mechanics in reproducing protein conformational distributions in molecular dynamics simulation. J Comput Chem. 2004. https://doi.org/10.1002/jcc.20065

23. Yang M, Huang J, Simon R, Wang LX, MacKerell AD, Jr. Conformational Heterogeneity of the HIV Envelope Glycan Shield. Sci Rep. 2017;7(1):4435. https://doi.org/10.1038/s41598-017-04532-9 PMID:28667249

24. Yang M, Simon R, MacKerell AD. Conformational Preference of Serogroup B Salmonella O Polysaccharide in Presence and Absence of the Monoclonal Antibody Se155-4. J Phys Chem B. 2017;121(15):3412–23. https://doi.org/10.1021/acs.jpcb.6b08955

25. Wang L, Friesner RA, Berne BJ. Replica Exchange with Solute Scaling: A More Efficient Version of Replica Exchange with Solute Tempering (REST2). J Phys Chem B. 2011;115(30):9431–8. https://doi.org/10.1021/jp204407d

26. Liu P, Kim B, Friesner RA, Berne BJ. Replica exchange with solute tempering: A method for sampling biological systems in explicit water. Proc Natl Acad Sci U S A. 2005;102(39):13749.

27. Hlozek J, Kuttel MM, Ravenscroft N. Conformations of Neisseria meningitidis serogroup A and X polysaccharides: The effects of chain length and O-acetylation. Carbohydr Res. 2018;465:44–51. https://doi.org/https://doi.org/10.1016/j.carres.2018.06.007

28. Hsieh PF, Wu MC, Yang FL, Chen CT, Lou TC, Chen YY, et al. D-galactan II is an immunodominant antigen in O1 lipopolysaccharide and affects virulence in Klebsiella pneumoniae: implication in vaccine design. Front Microbiol. 2014;5:608. https://doi.org/10.3389/fmicb.2014.00608 PMID:25477867

29. Guvench O, Greene SN, Kamath G, Brady JW, Venable RM, Pastor RW, et al. Additive empirical force field for hexopyranose monosaccharides. J Comput Chem. 2008;29(15):2543–64. https://doi.org/10.1002/jcc.21004 PMID:18470966

30. Raman EP, Guvench O, MacKerell AD. CHARMM Additive All-Atom Force Field for Glycosidic Linkages in Carbohydrates Involving Furanoses. J Phys Chem B. 2010;114(40):12981–94. https://doi.org/10.1021/jp105758h

31. Jorgensen WL, Chandrasekhar J, Madura JD, Impey RW, Klein ML. Comparison of simple potential functions for simulating liquid water. J Chem Phys. 1983;79(2):926–36.

32. Brooks BR, Brooks III CL, MacKerell AD, Jr., Nilsson L, Petrella RJ, Roux B, et al. CHARMM: the biomolecular simulation program. J Comput Chem. 2009;30(10):1545–614. PMID:19444816

33. Brooks BR, Bruccoleri RE, Olafson BD, States DJ, Swaminathan S, Karplus M. CHARMM: A program for macromolecular energy, minimization, and dynamics calculations. J Comput Chem. 1983;4(2):187–217. https://doi.org/10.1002/jcc.540040211

34. MacKerell AD, Brooks B, Brooks CL, Nilsson L, Roux B, Won Y, et al. CHARMM: The Energy Function and Its Parameterization. Encyclopedia of Computational Chemistry: John Wiley & Sons, Ltd; 2002.

35. Hoover WG. Canonical dynamics: Equilibrium phase-space distributions. Phys Rev A. 1985;31(3):1695–7. https://doi.org/10.1103/PhysRevA.31.1695

36. Feller SE, Zhang Y, Pastor RW, Brooks BR. Constant pressure molecular dynamics simulation: The Langevin piston method. J Chem Phys. 1995. https://doi.org/10.1063/L470648

37. Ryckaert J-P, CIccotti G, Berendsen HJC. Numerical integration of the cartesian equations of motion of a system with constraints: molecular dynamics of n-alkanes. J Comput Phys. 1977;23(327-341).

38. Darden T, York D, Pedersen L. Particle mesh Ewald: An N·log(N) method for Ewald sums in large systems. J Chem Phys. 1993;98(12):10089–92. https://doi.org/10.1063/L464397

39. Eastman P, Friedrichs MS, Chodera JD, Radmer RJ, Bruns CM, Ku JP, et al. OpenMM 4: A Reusable, Extensible, Hardware Independent Library for High Performance Molecular Simulation. J Chem Theory Comput. 2013;9(1):461–9. https://doi.org/10.1021/ct300857j PMID:23316124

40. Jiang W, Hodoscek M, Roux B. Computation of Absolute Hydration and Binding Free Energy with Free Energy Perturbation Distributed Replica-Exchange Molecular Dynamics (FEP/REMD). J Chem Theory Comput. 2009;5(10):2583–8. https://doi.org/10.1021/ct900223z PMID:21857812

41. Yang M, MacKerell AD, Jr. Conformational sampling of oligosaccharides using Hamiltonian replica exchange with two-dimensional dihedral biasing potentials and the weighted histogram analysis method (WHAM). J Chem Theory Comput. 2015;11(2):788–99. https://doi.org/10.1021/ct500993h PMID:25705140

42. Brooks BR, Brooks CL, Mackerell AD, Nilsson L, Petrella RJ, Roux B, et al. CHARMM: The biomolecular simulation program. J Comput Chem. 2009;30(10):1545–614. https://doi.org/10.1002/jcc.21287

43. Wodak SJ, Janin J. Analytical approximation to the accessible surface area of proteins. Proc Natl Acad Sci U S A. 1980;77(4):1736–40. PMID:16592793

44. Hasel W, Hendrickson TF, Still WC. A rapid approximation to the solvent accessible surface areas of atoms. Tetrahedron Comput Methodol. 1988;1(2):103–16. https://doi.org/https://doi.org/10.1016/0898-5529(88)90015-2

45. Humphrey W, Dalke A, Schulten K. VMD: visual molecular dynamics. J Mol Graph. 1996;14(1):33–8, 27-8. PMID:8744570

46. Baliban SM, Yang M, Ramachandran G, Curtis B, Shridhar S, Laufer RS, et al. Development of a glycoconjugate vaccine to prevent invasive Salmonella Typhimurium infections in sub-Saharan Africa. PLoS Negl Trop Dis. 2017. https://doi.org/10.1371/journal.pntd.0005493

